# Bacteriome diversity of blackflies gut and association with *Onchocerca volvulus*, the causative agent of onchocerciasis in Mbam valley (Center Region, Cameroon)

**DOI:** 10.1101/2020.06.12.148510

**Authors:** Arnauld Efon-Ekangouo, Hugues Nana-Djeunga, Guilhem Sempere, Joseph Kamgno, Flobert Njiokou, Paul Moundipa-Fewou, Anne Geiger

## Abstract

**Background:** Vector control using larvicides is the main alternative strategy to address limits of preventive chemotherapy using ivermectin to fight onchocerciasis. However, it remains substantially limited by implementation difficulties, ecological concerns and resistance of vector populations. Therefore, efficient and environmentally safe alternative control strategies are still needed. This study explores the role of blackfly bacterial communities both on vector competence and refractoriness to *O. volvulus* infection in order to determine their potential as a novel vector control-based approach to fight onchocerciasis.

**Principal findings:** A total of 1,270 blackflies were dissected and the infection rate was 10.1%, indicative of ongoing transmission of onchocerciasis in the surveyed communities. Sequencing process revealed 19 phyla and 210 genera, highlighting the diversity of gut blackflies bacterial communities. *Wolbachia* was the predominant genus with 70% of relative abundance of blackflies gut bacterial communities. *Serratia sp* and *Acidomonas* genera were significantly abundant among infected blackflies (p=0.043 and p=0.027, respectively), whereas other genera as *Brevibacterium* were associated with the absence of infection (p=0.008).

**Conclusion/Significance:** This study revealed that blackfly native bacteria are potentially involved in infection by *O. volvulus*, either by facilitating or preventing the parasite infestation of the vector. These bacteria represent an interesting potential as a biological target for a novel approach of vector control to fight onchocerciasis.

**Author summary:** Studies of arthropods involved in vector-borne diseases (tsetse flies, mosquitoes, and drosophila) demonstrated the importance of their native bacteria either to ease infection and transmission of human pathogenic microorganisms including parasites or on the contrary to induce host protective effects against these parasites. Indeed, some native bacteria of arthropod vectors are now recognized to be associated either with the resistance of their hosts to parasitic infections, or the reduction of their host’s viability in case of the parasite infestation, thus highlighting the potential of such bacteria to be used as biological tool for vector control strategies. However, such bacteria have never been described on blackfly, an arthropod transmitting *Onchocerca volvulus*, which is the parasite responsible of onchocerciasis commonly known as river blindness. This study aimed to fill this gap by investigating the bacterial diversity of blackfly bacteriome and describing the possible role of bacteria communities in susceptibility/resistance features of the blackflies to *O. volvulus* infection, and therefore their potential as biological targets or tool for vector control. The screening of these blackflies’ native bacteria during this study, highlighted some bacteria genera of interest with significant association either with the absence of *O. volvulus* in blackfly or with vector infection.

## Introduction

Onchocerciasis or river blindness is an infectious disease caused by the parasitic nematode, *Onchocerca volvulus*. The vector, a blackfly of the genus *Simulium*, is an arthropod that breeds in the oxygenated waters of fast flowing rivers [1]. Following ingestion of microfilariae by the vector during its blood meal, the first stage larva penetrates the midgut wall and migrates to the fly muscles where it molts twice. The third stage larva migrates to the head of the blackfly [1,2] and penetrates the skin of human, the only known natural vertebrate host of *O. volvulus*, during a subsequent blood meal. The larvae migrate to the subcutaneous tissue where they form nodules and reach adult stage, with an average lifespan estimated to 10-15 years [2,3]. After maturation and mating, adult females will release 200,000 - 400,000 microfilariae per three monthly reproductive cycle for their entire life. Microfilariae may then invade the dermis causing skin conditions, as well as eye tissues causing various eye lesions (keratitis, iridocyclitis…) which ultimately result in permanent blindness [4]. Indeed, onchocerciasis is the second cause of blindness of infectious origin after trachoma [5].

Approximately 120 million people are at risk of contracting the disease worldwide, and 37 million are believed to be infected [6,7]. Africa has the highest burden of the disease, with 99% of the infection and 1.49 million disability-adjusted life years (DALYs) annually. It has been reported to be significantly associated with, epilepsy [8,9] and excess mortality [10,11] among people living in endemic areas.

Ivermectin, the only safe and effective anthelmintic with microfilaricidal effect on *O. volvulus*, was registered for the control of onchocerciasis in 1987 [12,13]. Preventive chemotherapies through the community-directed treatment with ivermectin (CDTI) strategy led to the interruption of the transmission of the disease in four of the six onchocerciasis foci in Latin America [14,15]. However, despite almost three decades of preventive chemotherapy in Africa, onchocerciasis remains a public health problem in many countries, including Cameroon [6,16]. Indeed, recent epidemiological surveys carried out between 2011 and 2015, revealed the persistence of onchocerciasis with microfilarial prevalence higher than 60% in certain foci in the Centre, Littoral and West Regions despite more than two decades of CDTI [6,16]. The reasons related to this situation appear to be multifactorial, including (i) high proportion of permanent non-compliant infected persons living in endemic areas [17,18] (ii) foci located in conflict and hard to reach zones [12], (iii) sub-optimal responses of *Onchocerca volvulus* to ivermectin [19-22] (iv) very high transmission levels due to high densities of black flies with important vector competence [23]. These factors constitute tremendous obstacles to the process of elimination of the disease [20,24].

In order to accelerate the interruption of transmission process, various complements or alternatives to the classical CDTI approach, so-called alternative/complementary treatment strategies, have been considered, including vector control [25-27]. However, the classical vector control approach based on the weekly use of larvicides, either aerial or ground larviciding in blackflies infested breeding sites, remains limited by the implementation difficulties, the significant risks of ecological pollution and fairly substantial implementation costs [26] and foci specificities constraints related to the geography and the size of rivers which are substantially important [17,28]. Also, re-colonization of blackflies after treatment of breeding sites has been observed in some foci.

These vector control difficulties being shared by other vector-borne diseases, mitigation or alternative approaches are likely to be the same. Previous studies in other vectors (tsetse flies, mosquitoes, and *Drosophila*) demonstrated the impact of their microbiome in the vector competence, as well as their promising role as effective tools/targets for new generations of vector control strategies [29-31]. It is now well known that certain native bacteria species such as *Wigglesworthia glossinidia* an obligate intracellular bacteria of tsetse intestinal cells, are necessary for the survival of their host [32,33]; While some bacteria are associated with the refractory character of their hosts to parasitic infection, others are associated with the reduction of the viability of their hosts in case of the parasite infestation. This evidence is observed with *Serratia mascesens* which produces a trypanolytic compound preventing the establishment of *Trypanosoma cruzi* in the digestive tract of *Rhodnius prolixus* [33,34], although other *Serratia* species have been associated with the reduction of *Anopheles* infection by *Plasmodium* [35].

Hence, native bacteria of vectors can be targeted and/or manipulated in different ways for vector control, notably as chemotherapeutic target, immunological reinforcement or cytoplasmic incompatibility, by inducing through genetic manipulation, a disturbance of molecular interactions between the parasite and vector host [31,36]. The prerequisite for the development of such vector control approach is the identification and the characterization of native bacterial communities of the targeted vector and the assessment of their potential as effective tool/target for vector control. Thus, the discovery of such bacteria in blackflies would constitute a major breakthrough and will open wide avenues for the development of an innovative approach in the fight against onchocerciasis in Africa through the development of non-infestable blackflies. Hence, this study was designed to screen the whole bacterial communities of blackfly gut and highlight bacterial species associated with vector competence and those associated with vector refractoriness to *O. volvulus* infection and thus assess their possible impact on onchocerciasis transmission.

## Methods

### Ethics approval and consent to participate

Although this study did not directly involve human subjects, sample (capture of blackflies) were collected using the human landing collection technique which require volunteers. Hence, an ethical clearance was obtained from the Centre Regional Ethics Committee for Human Health Research (N°1011/CRERSH/C/2020) and administrative authorizations were granted by the Centre Regional Delegate for Public Health and the Bafia District Medical Officer. Prior to the beginning of the entomological survey, the objectives and schedules of the study were explained to all the volunteers. Participation was entirely voluntary and each of them (aged 24 years and above) was free to opt out without fear of retaliation. The volunteers recruited lived in sampling sites, so they were not more exposed to fly bites than usual. Moreover, volunteers were trained to capture flies before being bitten. Finally, ivermectin was provided as preventive chemotherapy against onchocerciasis.

### Study area

This study was carried out in the Bafia health district, situated in Mbam and Inoubou Division, Centre Region, Cameroon. This health district is known for its historical endemicity to onchocerciasis and disease persistence despite two decades of ivermectin-based preventive chemotherapy. Communities of this health district are mainly watered by the Mbam River and its tributaries whose falls and rapids promote and maintain throughout the year blackfly breeding sites. The phytogeography of this area shows a forest/savannah transition zone dominated by a peri-forest savannah with forest galleries along the rivers and important breeding sites favorable to the development of blackflies. Bafia is mainly dominated by the subequatorial climate with average temperature of 23.5°C and bimodal rainfall regime marked by modest precipitations with average rainfall of 831.7 mm. Socio-economic activities are dominated by sand extraction in the Mbam river, as well as agriculture and trade on the shores of the latter.

### Capture, dissection and preservation of blackflies

Entomological surveys were conducted on April 2019 in three communities of the Bafia health district, namely Bayomen (04° 51’52’’N; 011° 06’07’’E), Biatsota (04° 41’11’’N; 011° 17’28’’E) and Nyamongo (04° 46’57’’N; 011° 17’24’’E). In each selected site, blackflies were captured using the “human Landing collection” method. Indeed, catches were made up by two groups of volunteers, the first working from 8 a.m. until 1 p.m. and the second from 1 p.m. until 5 p.m. Only female blackflies, which are hematophagous, landed on exposed legs of well-trained community volunteers, and captured before having time to take their blood meal. Captured blackflies were individually dissected in situ for parity, under sterile conditions using a binocular magnifier. For each identified parous blackfly, gut, thorax, head and feet were separated and transferred individually into well labelled 1.5 mL Eppendorf tube containing 70° Ethanol and stored at -20°C for further molecular analysis.

### DNA extraction and *O. volvulus* PCR amplification

Genomic DNA was extracted from separated parts of fly (head, thorax, gut and feet) and purified on MiniElute PCR purification columns using the QIAamp DNA Mini kit (Qiagen Inc., Les Ulis, France) and eluted in 50 µL molecular biology-grade water. DNA samples from thorax and feet were stored for further purposes. DNA extracted from gut and head samples were used for the detection of *O. volvulus* by quantitative PCR (qPCR) using specific primers (Forward: 5’-GCTATTGGTAGGGGTTTGCAT-3’ and reverse: 5’-CCACGATAATCCTGTTGACCA-3’) targeting a DNA portion (128bp) of ND5 *O. volvulus* gene and probe (5’-FAM-TAAGAGGTTAAGATGG NFQ-3’). Each well of the microtiter plate (MicroAmp fast optical 96-well reaction plate, Applied Biosystems) was filled with 20 µL of final solution, containing 2 µL DNA template and 18 µL of PCR master mix made up of: 12 µL molecular biology-grade water, 2 µL of 10× PCR buffer, 2.4 µL 50× MgCl_2_ (50 mM), 0.1 µL dNTPs (10 mM), 0.6 µL forward primer (10 mM), 0.6 µL reverse primer (10 mM), 0.2 µL ND5 *O. volvulus* probe, and 0.1 µL HotStarq polymerase (5 U/µL). For each amplification process, negative and positive controls were used to ensure good interpretation of final results. Real-time PCR assays were performed on an Applied Biosystems Step One Plus real-time PCR machine (Applied Biosystems, Foster City, CA, USA) using the following cycling conditions: Initial denaturation at 95°C for 10 min, followed by 45 cycles, each including a denaturation step at 95°C for 1min, an annealing and elongation step at 60.1°C during 30s.

### High throughput sequencing and Meta-barcoding analysis

The 16S rRNA gene V3-V4 variable region was amplified using specific primers designed in the scope of a previous study [33] to assess the bacterial communities of blackfly guts using the Illumina MiSeq sequencing approach (MR DNA Laboratory, http://www.mrdnalab.com/shallowater, USA). PCR was performed using the HotStar Taq Plus Master Mix Kit (Qiagen Inc, Texas, USA) under the following conditions: 94°C for 3 min for initial denaturation, followed by 30 cycles of successive steps: denaturation at 94°C for 30 s, annealing at 53°C for 40 s and elongation at 72°C for 1 minute, and a final elongation step at 72°C for 5 min. After amplification, PCR products were checked on 2% agarose gel to determine the success of amplification and the relative intensity of bands. Multiple PCR products were pooled together in same proportions based on their molecular weight and DNA concentrations. Pooled PCR products were purified using calibrated Ampure XP beads (Details on Manufacturer). Then the pooled and purified PCR product were used to prepare Illumina DNA library. Sequencing process was performed at MR DNA (www.mrdnalab.com, Shallowater, TX, USA) using a MiSeq following the manufacturer’s guidelines.

Prior to running the metabarcoding pipeline, a specific reference file for the assignment step was generated. This was achieved by running CutAdapt v1.8 (http://dx.doi.org/10.14806/ej.17.1.200) with those primers to extract V3-V4 reference sequences from the SILVA SSU database (release 132). The generated sequences were deposited in the EMBL-EBI (Study accession number: PRJEB38276; secondary study accession number: ERP121684).

The first stage in the workflow consisted in filtering read quality using CutAdapt with a threshold value of 20. Then, VSearch v2.14 (https://dx.doi.org/10.7717%2Fpeerj.2584) was used in combination with CutAdapt for the following series of tasks: (1) merging the forward and reverse reads of each sample; (ii) demultiplexing to obtain one fastq file per sample; (iii) clipping barcodes and primers; (iv) excluding sequences containing unknown bases; (v) calculating expected error rate; and (vi) performing sample-level dereplication. The remaining sequences were then pooled into a single FASTA file to allow VSearch to carry out a global dereplication, after which clustering was applied to remaining sequences using Swarm v2.2.2 (https://dx.doi.org/10.7717%2Fpeerj.1420). VSearch was then used again to identify chimeric clusters.

The STAMPA (https://github.com/frederic-mahe/stampa) pipeline was then run for taxonomic assignment of representative OTU sequences based on the contents of the specific reference file generated from SILVA SSU records. This generated an OTU table to which the following filters were applied in order to retain targeted taxa at genus level: elimination of clusters with a high expected error (>0.0002), elimination of small clusters (less than 3 sequences) observed in a single sample.

Data obtained after analysis of raw data from sequencing have been analyzed using Calypso 8.1 (https://dx.doi.org/10.1093%2Fbioinformatics%2Fbtw725), an online software dedicated to bacterial taxonomic analysis. Rarefaction curves assessing the sequencing depth of each sample were performed prior to quantitative (assessing alpha and beta diversity) and comparative analyses between infected and uninfected flies, and sampling sites. Significant differences in bacterial richness between infected and uninfected flies, and between the three sampling sites were tested using nonparametric Kruskal-Wallis test with threshold of significance set at α= 0.05.

## Results

### Blackflies infection by *Onchocerca volvulus*

From the 1,270 blackflies caught and dissected, a total of 207 (16.3%) were parous and analyzed for infection (Table 1). Overall, 21 (10.1%) blackfly guts were infected with *O. volvulus*. The infection rate was capture site dependent. Indeed, Biatsota displayed the highest infection rate (13.0%), followed by Bayomen (8.8%) and Nyamongo (6.8%), though the difference was not statistically significant. On DNA samples from blackfly heads, only 5 samples were confirmed positive to *O. volvulus* detection, with an infectivity rate of 2.4%, and differences in infectious blackflies distribution among geographic areas was not significant.

**Table 1.** Distribution of collected blackflies according to geographic origin.

| Geographic origin | No. Dissected flies | No. Parous flies | No. dissected flies with *O. volvulus* (%) | No. parous flies with *O. volvulus* (%) |
| --- | --- | --- | --- | --- |
| Bayomen | 200 | 34 | 3 (1.5) | 3 (8.8) |
| Biatsota | 558 | 100 | 13 (2.3) | 13 (13.0) |
| Nyamongo | 512 | 73 | 5 (1.0) | 5 (6.8) |
| **Total** | **1270** | **207** | **21 (1.7)** | **21 (10.1)** |

No.: number of; *O. volvulus*: *Onchocerca volvulus*

### Sequencing data analysis

DNA from 42 blackflies (21 infected and 21 randomly chosen among uninfected), were selected for sequencing. Sequencing of 16S rDNA from total DNA extracted from blackflies intestine using Illumina sequencing technology generated a total of 3,427,049 high quality sequence reads across the V3-V4 region. The average number of tags per sample for the V3–V4 region was 81,596 (ranging from 11,351 to 136,972 per sample), with read length varying from 300 to 400 nucleotides. The sequencing depth was performed to assess how well sequence data represent the diversity of the studied microbial communities. Consequently, a rarefaction curve (Fig 1) showed the saturation of most of them between 60,000 and 100,000 reads, indicating that the mean sequencing effort was sufficient to characterize almost all the OTUs. Nonetheless, some samples showed poorly amplified OTUs, in particular the reference samples AP10, AN40 and AN35.

**Fig 1.**
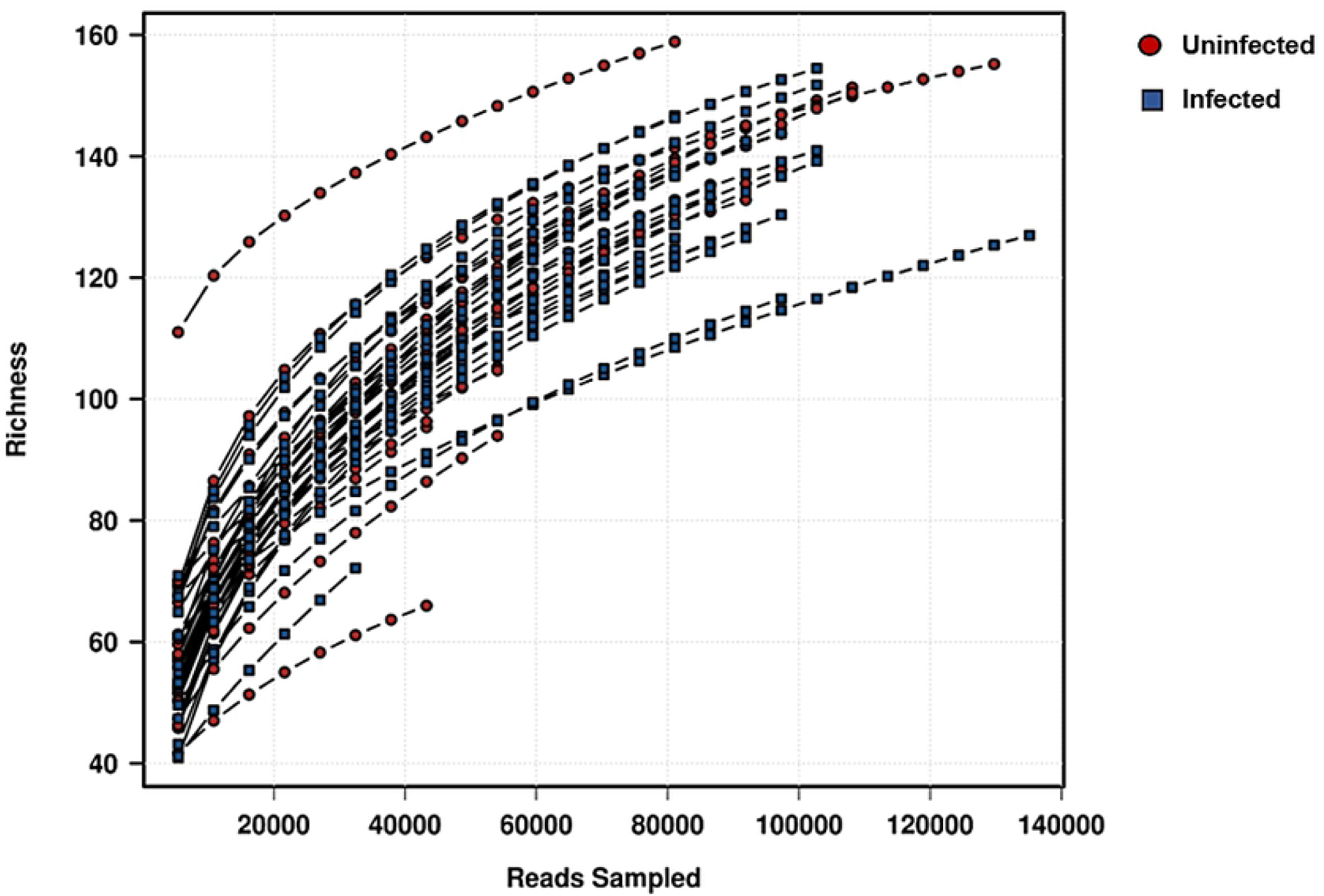
Rarefaction analysis on the blackfly samples.

### Taxonomic Assignation

The taxonomic assignation of OTUs sequences allowed to identify 23 phyla among which 22 belonged to the Bacteria kingdom and only one belongs to *Archaea (Euryarchaeota* phylum). For further analysis, we removed rare taxa by excluding those presenting with less than 0.01% of relative abundance across all samples and those present in less than 10% of sequenced samples to exclude potential contaminants. Using these filters, a total of 19 phyla were found fulfilling stated criteria and retained for further investigations. Among retained phyla, *Proteobacteria, Unclassified bacteria* and *firmicutes* represented the most predominant Bacteria phyla with 44.7%, 15.4% and 9.6% of mean relative abundance across the 42 samples, respectively. *Proteobacteria* was the most important phylum (>90%) in 22 out of 42 samples. This was confirmed by the heat map (Fig 2) showing the distribution of the mean relative abundance of the 19 retained phyla, and highlighting the predominance of *Proteobacteria*, significantly found in almost all samples. Other phyla were unevenly distributed and abundant among the different samples. In fact, *Caldiserica, Kiritimatiellaeota*, and *Thermotogae* were the less represented phyla among the analyzed samples, found only in four samples with non-significant relative abundance within all samples. NoHit phylum represents taxonomic description whose sequences do not fit with any OTU listed in the databases. The hierarchical clustering of phyla relative abundance distinctly shows three clusters namely cluster A including samples from AP15 to AN29, cluster B from AP11 to AN24 and cluster C from AN27 to AN34. Moreover, cluster B, made up of three sub-trees was organized in two distinct groups based on the distribution of *Firmicutes, Actinobacteria* and *Bacteriodetes* which were substantially abundant in cluster B1 (AN16 to AN24) than in cluster B2 (AP11 to AN38). The clustering does not however match with specific condition, either geographic or infection status.

**Fig 2.**
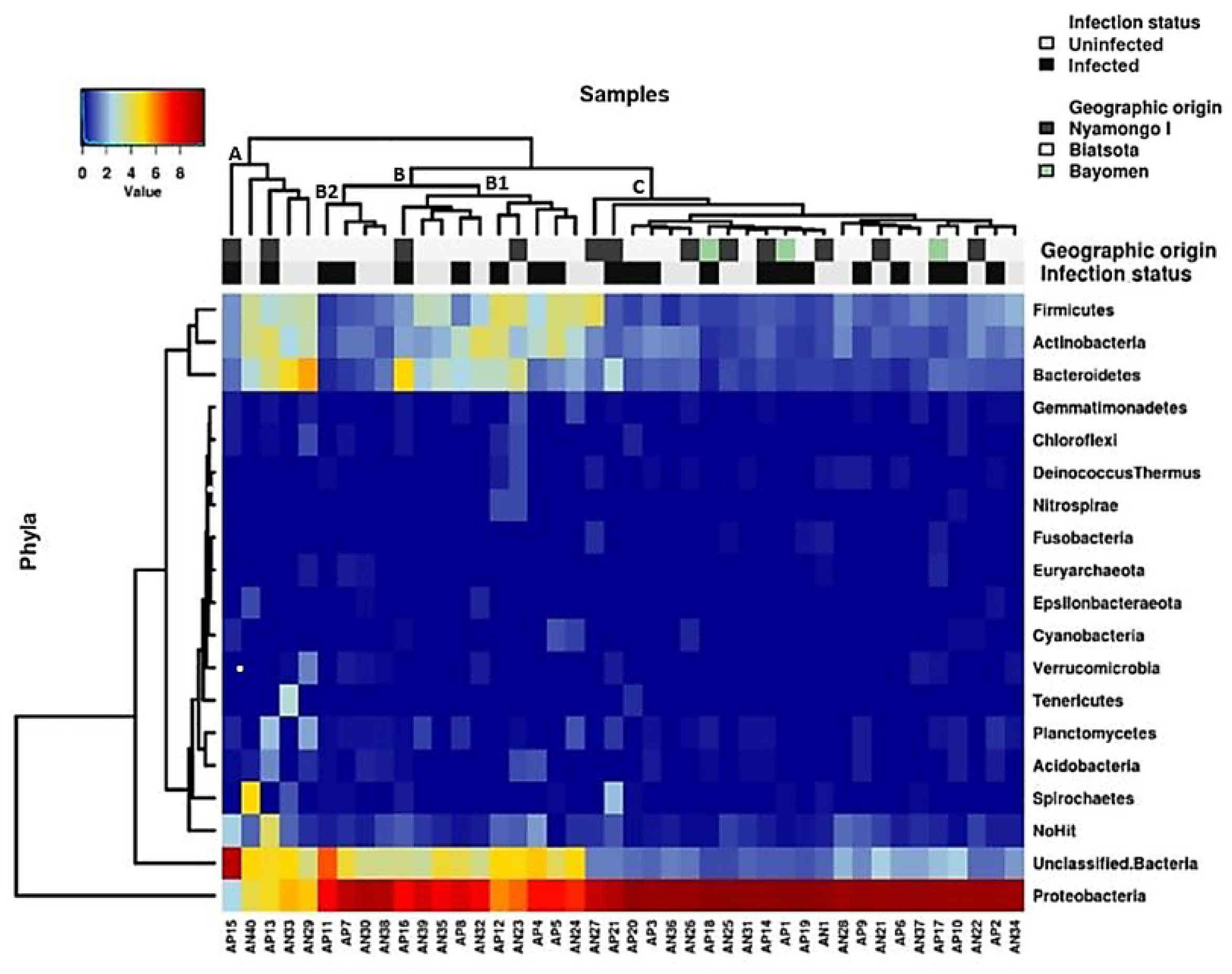
Heat map analysis of the distribution and abundance of the bacterial phyla in blackfly gut samples. Clusters are organized from the left to right: cluster A including samples from AP15 to AN29, cluster B from including samples AP11 to AN24 and cluster C including samples from AN27 to AN34. The cluster B is divided into two sub-cluster: Sub-cluster B1 including samples from AN16 to AN24 and sub-cluster B2 including samples from AP11 to AN38.

### Bacterial genera

A total of 554 bacterial genera was observed, with relative abundance ranging from 6.58619E-05% to 70.16%. Likewise phyla analysis, we excluded bacterial genera with relative abundance lower than 0.01%, and those present in less than 10% of samples to exclude potential contaminants, and a total of 210 genera were thus retained for further analysis.

Twenty bacterial species were shown to be systematically present in all samples with various relative abundance. The bacterial taxa could possibly represent the blackfly gut core microbiota. A heat map analysis (Fig 3) showed that the hierarchical clustering of the bacterial relative abundance of these 20 bacterial genera across the 42 samples resulted in a poorly structured tree. Only *Wolbachia* was distinctly separated from the other genera of which abundance was unevenly distributed among the samples. However, the map shows a slight contrast on these bacterial distribution with a more uniform color print at the right half of the heat map. This observation was strengthened by the hierarchical clustering of the samples on the basis of the relative abundance of considered bacterial genera, which allowed discriminating two main clusters: cluster 1 including samples AP17 to AN23, and cluster 2 including AP16 to AN31 (Cluster’s indicative are shown on Fig 3). However, the samples of either cluster 1 or cluster 2 seem to be associated neither with sample infection status nor with sample geographic origin. The hierarchical clustering of bacteria genera based on blackflies and infection status and geographic area where they were captured, showed no specific bacterial clustering. However, *Wolbachia* genus showed a relative homogeneity of abundance distribution from samples AP16 to AN31. This observation matches with the cluster 2 identified in the preceding hierarchical clustering description (S1 Table).

**Fig 3.**
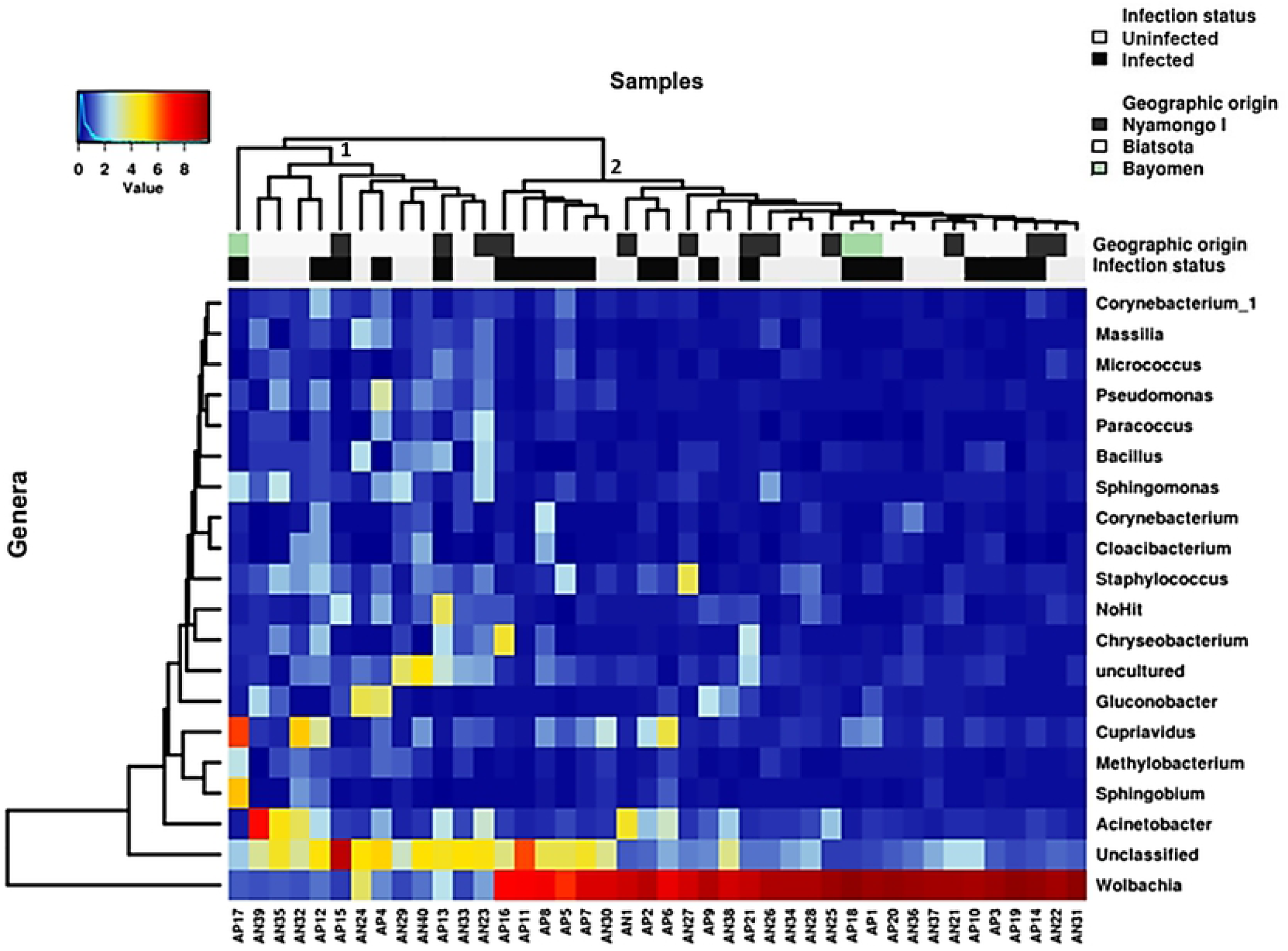
Heat map analysis of the distribution and abundance of the bacterial genera in blackfly gut samples. Samples are clustered from left to right: cluster 1 including samples AP17 to AN23 and cluster 2 including samples AP16 to AN31.

The most prominent bacteria was the genus *Wolbachia* with 70.16% of relative abundance (1.3 and 94.1% of relative abundance among samples) (Fig 4A), followed by *Gluconobacter* and *Acinetobacter* genera, with 4.0% (0.1 and 46.8) and 3.5% (0.2 and 59.5) of relative richness among OTUs, respectively. The less abundant bacterial genera were *Gemmatirosa* and *Modestobacter*, with a relative richness of 0.0102 and 0.01, respectively. This relative abundance distribution, largely occulted by *Wolbachia* genus, is substantially modified after exclusion of this genus, with the evidence of other bacterial genera (Fig 4B).

**Fig 4.**
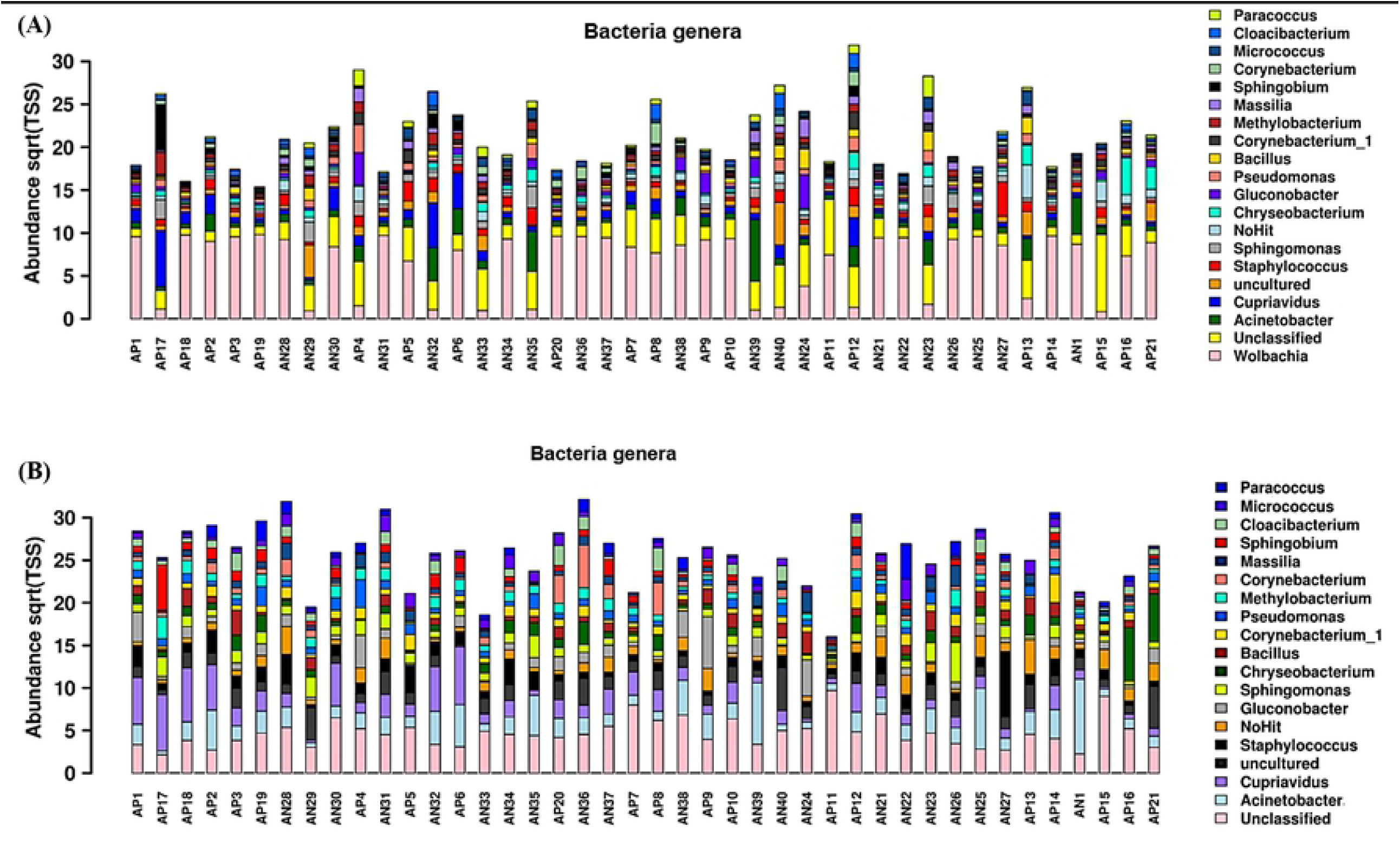
Relative abundance of the bacterial genera along the 42 blackflies samples: (A) All the 20 bacteria genera including *Wolbachia*. (B) All 19 bacteria genera, without *Wolbachia* genus.

### Association between blackflies’ bacterial diversity with either their infection status or geographical origin

We investigated the potential relationship between the bacterial diversity of blackflies gut, with either the infection status or geographical origin by estimating the α-diversity using Shannon Index which measures overall diversity, including both the number of OTUs and their evenness. No significantly differences were observed both on bacterial diversity and richness regarding the geographical origin (p = 0.387) (Fig 5A) and the infection status (infected vs. non-infected) of sampled blackflies (p = 0.349) (Fig 5B).

**Fig 5.**
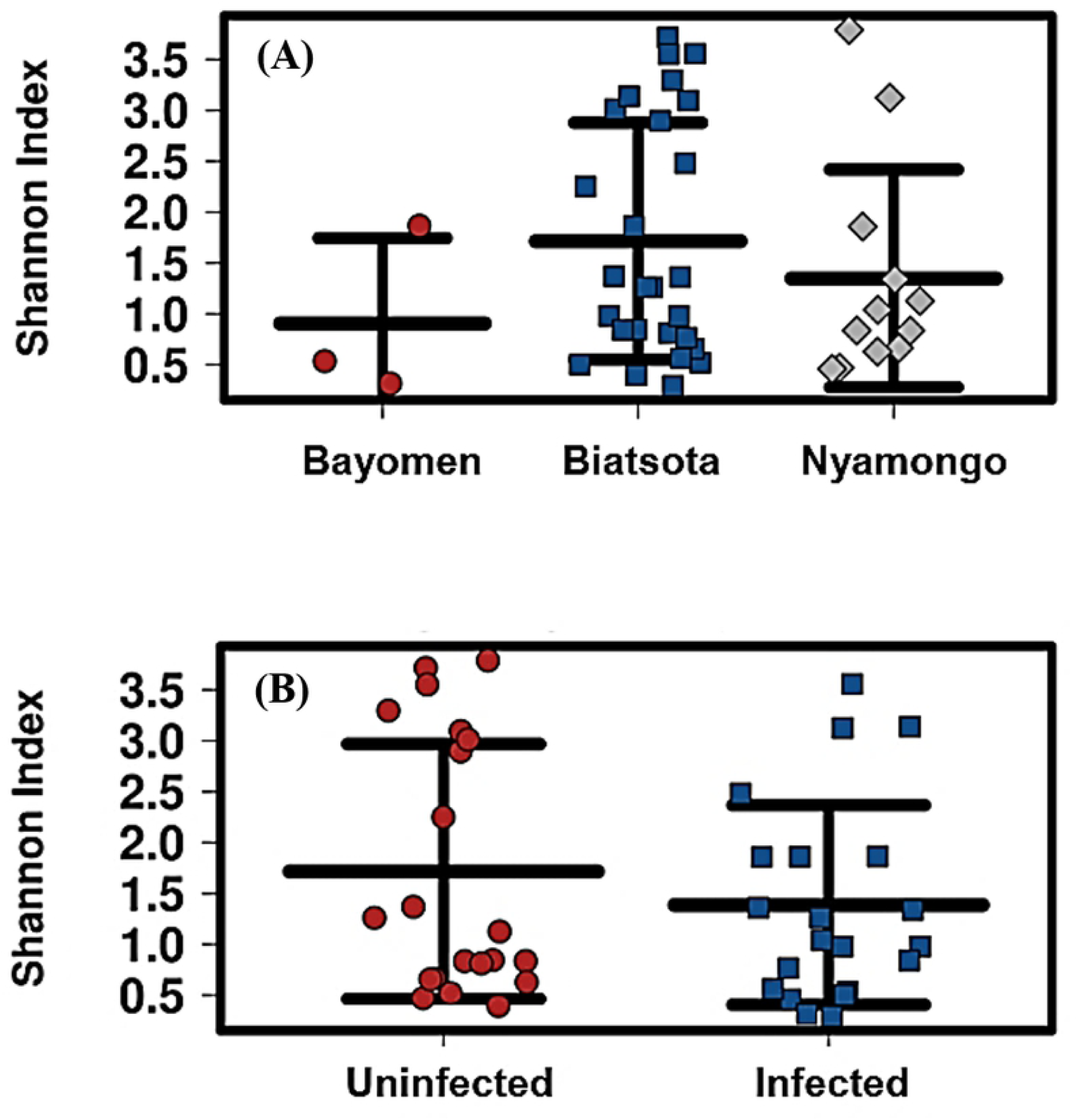
Alpha diversity using Shannon Index assessing the relationship between the bacterial diversity of blackflies gut: (A) with geographical origin, and (B) with blackfly infection status. The same non significance has been observed when using other metrics (evenness, richness and Simpson Index).

### Multivariate association between bacterial diversity in blackflies, infection status and geographical origin

The investigation of potential relationships between the tripartite factors: structure of gut bacterial communities of blackflies, their geographical origin (Bayomen, Biatsota and Nyamongo) and their infection status (infected vs. uninfected) was done to highlight potential complex associations between gut blackfly bacterial composition and the two other covariates. A hierarchical clustering using the Bray–Curtis index did not discriminate unambiguously the different groups with regard to sample origins or infection status (Fig 6). However, it showed a structure quite similar (symmetrical from the point of view of the layout) to the one on heat map (Fig 3), with 2 main clusters, cluster I from A24 to AP8 and cluster II from AN35 to AP17. The composition of cluster I is identical to the one of cluster 2 with two “curious” exceptions: the position of the sample AN29 is completely isolated in this tree while it was previously integrated in cluster 2, and sample AN24 previously located in cluster 1 (Fig 3) has shifted to cluster I now accounting for 30 samples.

**Fig 6.**
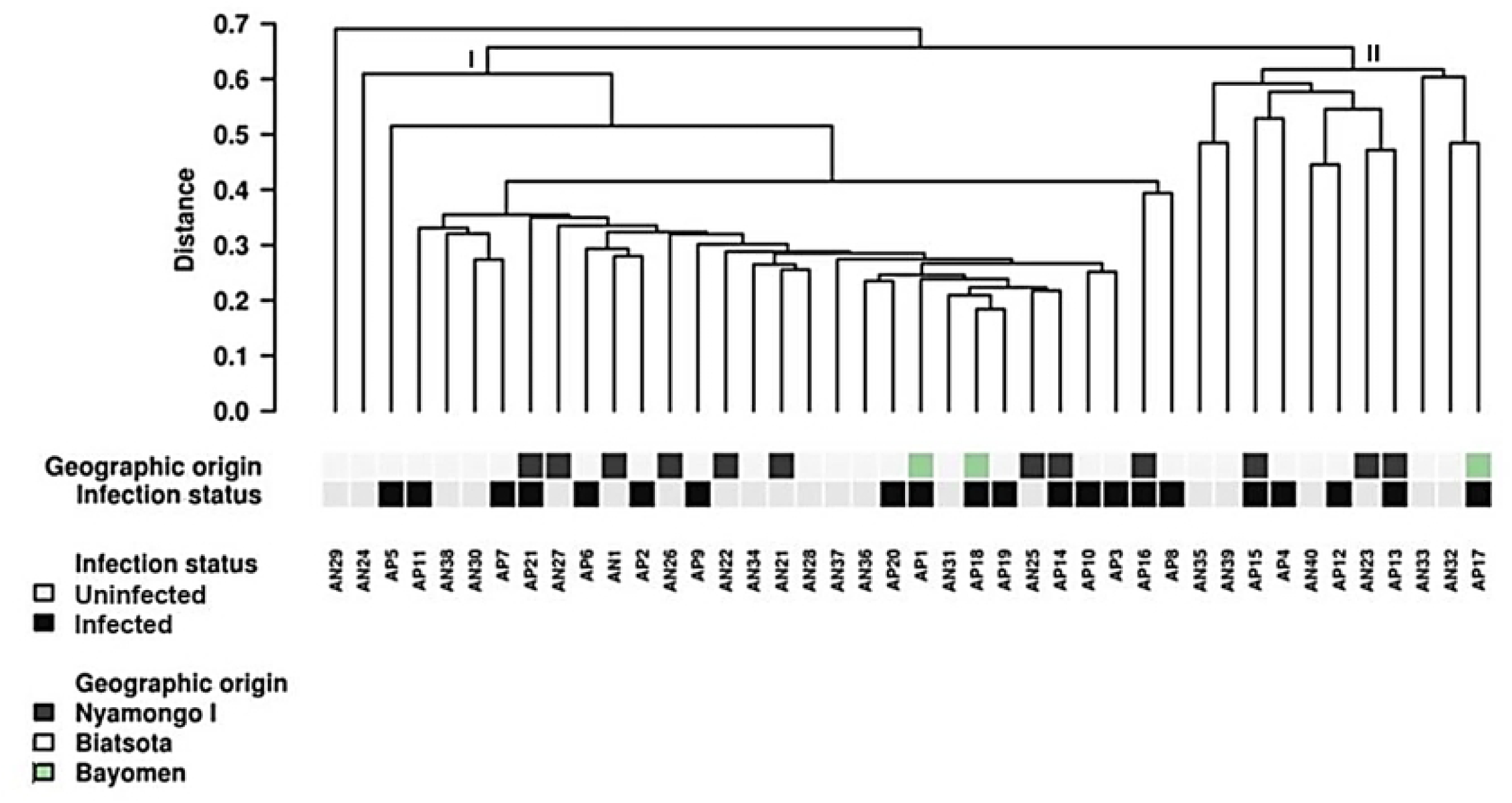
Bray-Curtis multivariate analysis of bacterial distribution across the 42 selected blackfly intestine samples. Two mains clusters are highlighted here: cluster I from A24 to AP8 and cluster II from AP35 to AP17

### Research of biomarker according to blackfly infection status

In order to determine potential biomarker(s) of bacterial community associated to specific geographical community and status infected/uninfected of sampled blackflies, we performed richness analysis of each bacterial genus in different conditions. This analysis that included the 100 most abundant bacterial genera within our samples using feature selection method (S1 Fig), allowed to identify five bacterial genera significantly associated to infected status, notably *Cyanothece_PCC7424* (p=0.032), *Serratia* (p=0.043), *Acidomonas* (p=0.027), *Roseamomas* (p=0.035) and *Cnuella* (p=0.046), whereas four bacterial genera were significantly associated to uninfected status, notably, *Sanguibacter* (p=0.048), *Fructobacillus* (p=0.00044), *Micrococcus*(p=0.034) and *Brevibacterium* (p=0.0087).

## Discussion

This study aiming to characterize the whole bacterial communities within the blackflies gut and the assessment of their potential associations with vector competence is, to our knowledge, a pioneer in onchocerciasis. Nonetheless, this approach based on successful outcomes of parallel studies on other vector-borne diseases is a fundamental prerequisite for application of vector control strategy-based on modified non-infestable blackflies to gradually reduce disease transmission in onchocerciasis endemic areas.

The global infestation rate was 10% in Bafia heath district, suggesting that onchocerciasis transmission is still ongoing in this historical endemic area despite almost three decades of community-directed treatment with ivermectin (CDTI). Our results confirm previous evidences supportive of ongoing transmission both within the human population (microfilarial prevalence varying from 24.4 to 57.0 %) [6] and in vector population (>98% of infection detected in blackflies using pool screening approach) [37]. The level of endemicity in surveyed communities of Bafia health district is related to their proximity with Mbam River, characterized by series of rapids and well oxygenated water that provide ideal breeding sites for blackfly. This observations are supportive of the importance of vector control approach in the process of elimination of onchocerciasis [27,38].

According to both infestation and infectivity rates, Biatsota was considered as the most active community in terms of disease transmission; this situation is likely to be related to its geographical situation which is closer to Mbam River (first line community) as compared to other surveyed communities. Moreover, in this community, economic activities are more intense along the river, thus increasing the biting rate in contrast to Bayomen and Nyamongo communities where inhabitants are living less closely to the river, hence contact with blackflies are less important.

In this study, taxonomic assignation allowed to identify a total 19 phyla and 210 genera (the most relevant with relative richness >0.01% and present in >10% samples). This result highlights the diversity of gut blackflies bacterial communities, which seems significantly larger than bacterial diversity reported in many recent studies on other arthropods of medical importance such as tsetse fly and *Aedes*. Indeed, studies on tsetse fly, using the same molecular approach showed that the gut bacteria communities where made up of 14 phyla and 83 different bacteria genera [32,33], meanwhile sequencing of 16S rRNA sequences in *Aedes* (vector of Dengue virus) using 453 pyrosequencing technique showed they were made up of six phyla of bacteria [39].

Our findings revealed that *Proteobacteria* was the predominant phylum with 77.1% of mean relative abundance. This predominant phylum common to several insects [32,33,36,40], plays a major role in energy management [41]. *Wolbachia* was the most important bacterial genus with 70.2% of mean relative abundance of blackflies gut bacterial communities. This result is in agreement with previous estimates suggesting that *Wolbachia* infects more than 65% of all insect species [42], though they are also widespread and common in other invertebrates such as arachnids, crustaceans, and nematodes [43,44]. Beyond their presence and their likely role in vector biology, *Wolbachia* also plays an important role in the development and pathogenesis of the main filarial parasites (*Onchocerca volvulus, Brugia malayi, Mansonella perstans* and *Wuchereria bancrofti)* [43,45,46] except *Loa loa* [47,48].

The analysis of bacterial taxa from blackflies gut did not show significant differences in bacterial composition on blackflies originating from the three surveyed communities. This could be explained by the fact that the three selected communities Bayomen, Bioatsota and Nyamongo are located in the same geographical area (Bafia health district), within∼50 km and hence share both the same bio-ecological (climate, flora) and environmental features. This observations was similar to those recorded on *Anopheles* [49] where no significant differences between the bacterial flora of the mosquitoes collected in similar ecological features foci in Cameroon. Such evidence was also observed with tsetse flies [32,33], demonstrating that bacterial composition of flies collected in Campo and Bipindi, two foci sharing similar ecological features, were not significantly different. In the line of these studies, vector populations from distinct geographical area with different eco-climatic features are expected to share significantly different bacterial communities. Such possibility was evidenced by Askoy et al [50] who reported differences in bacterial composition between distinct populations of tsetse flies transmitting *Trypanosoma rhodesiense*. However, differences could not be exclusively associated to the ecological differences of surveyed foci, but also with tsetse species (*G. fuscipes fuscipes, G. morsitans morsitans and G. pallidipes*) that are commonly found in different biotic and abiotic habitats. In this frame, even though *Simulium damnosum* complex is known as the important vector for *O. volvulus* in Cameroon [37,51,52], *S. yahense* and *S. squamosum* are associated with forest and forest-savannah transitional zones [37,52]. Further studies should be conducted on *Simulium* genetics of these localities to ensure if the highlighted homogeneity of bacterial communities within captured blackflies are shared by a common *Simulium* species or if there are different *Simulium* species with similar bacterial communities.

Similarly, the analysis of the abundance diversity of the 20 bacteria genera hosted by all the selected flies showed no significant differences, between *O. volvulus* infected or uninfected, blackflies. The possibility that some bacteria genera escaped molecular characterization may be considered. Indeed, results of the rarefaction curves allowed to expect almost all the OTUs to be characterized. However, if so, one may expect missing bacteria, if any, to be present in very low abundance. Besides, the overwhelming presence of the genus *Wolbachia* could lower the efficiency of amplification process of low abundant or rare bacteria genera with potential biological implications.

Nevertheless, when the 100 most represented genera (S1 Fig) are considered (which are not hosted by all the 42 selected flies), a significant association between the abundance of some of them (*Serratia, Acidomonas, Roseamomas* and *Cnuella, cyanothece_PCC7424*) and blackflies infection was evidenced. These bacteria potentially improve the susceptibility of blackflies to *Onchocerca volvulus* infection. Presently, only *Serratia* species has been described in other vector [32,34,53] and his role seems to be vector-dependent. In Mosquitoes, *Serratia odorifera* has been associated with the susceptibility of *Aedes aegypti* both to chikungunya virus [54] and dengue virus [55]. Meanwhile, other studies demonstrated the ability of *Serratia marcescens* to produce some trypanolytic compounds that increase the refractoriness of *Rhodnius prolixus* to *T. cruzi* infection [34]. Further investigations are therefore needed to identify *Serratia* species on blackflies, decipher the host-bacteria interactions as well as to assess whether the biological role is mediated by single bacteria species or by the whole significantly associated bacteria genera. These evidences illustrate the complexity of molecular interaction with biological impact on vector susceptibility or refractoriness to parasite infection according to the bacterial species. Besides, other bacteria genera, in particular *Brevibacterium*, were found significantly associated with the absence of infection among blackflies. This gram-negative bacterium was not yet reported to play a biological role in any vector-borne disease, thus opening a potential research avenue with possible outcome of interest.

In addition to these questions about the possible association between intestinal bacteria and the susceptibility / resistance of blackflies to infection with *O. volvulus*, a supplementary question is arising from the structure of the hierarchical clustering shown in Figs 3 and 6. The 42 samples are clearly distributed into two clusters that neither the geographic origin nor the infection status, can explain. Considering all these samples are coming from blackflies belonging to the same species (*Simulium damnosum complex*), one cannot incriminate a possible differentiation related to a species difference. The simple observation on the heat map highlights contrast in colors intensity which represents differences in abundance of the various bacteria and allows to discriminate the two clusters; this is even more evident when we consider *Wolbachia*. Hence, existence within the vector population, of a genetic diversity (existence of different genotypes) could be at the origin of the observed structuration. Thus it appears necessary to explore, besides the possible involvement of intestinal bacteria in blackflies infection, such an hypothesis in further investigation in order to get a better insight into the complex interactions between the three partners, the blackfly, its intestinal bacteria and the parasite that are together responsible for the transmission of onchocerciasis.

## Conclusion

This study exploring the blackfly bacteriome is to our knowledge a pioneer on onchocerciasis vector. It revealed that some bacteria genera are associated with the presence of *O. volvulus* in blackflies while others are refractory to it, giving an insight of biomarkers with interesting potential as biological tool/target for developing of non-infestable blackflies. However, this study had some limitations in such as vector speciation of analyzed blackflies as well as sequence depth that needed to be improve in some samples.

## Acknowledgements

The authors are thankful to Dr Philippe Nwane, M. André Domche and Mrs Guindoline Kweban for their valuable contributions during blackfly collection and dissection in the field, as well as to the human landing collection operators for their willingness to participate in this study. The authors are also grateful to Dr Jean Marc Tsagmo and M. François Sougal Ngambia Freitas for their guidance during bioinformatic analyses of data.

## Supporting information

**S1 Table. Summary of the overall sequencing raw data regarding the 42 blackfly samples**.

**S1 Fig. Forest plot illustrating the Odd ratio variation of top 100 bacterial genera (biomarker candidates) relative abundance depending on blackflies infection status (uninfected/infected)**.

